# Characterisation of the genomic sequence of an iflavirus, a protoambidensovirus, and of three microviruses detected in mosquitoes (*Aedes albopictus* and *Culex quinquefasciatus*)

**DOI:** 10.1101/2024.11.08.622592

**Authors:** Sarah François, Aymeric Antoine-Lorquin, Doriane Mutuel, Patrick Makoundou, Marco Perriat-Sanguinet, Sandra Unal, Hélène Sobry, Anne-Sophie Gosselin-Grenet, Mylène Ogliastro, Mathieu Sicard, Mylène Weill, Célestine Atyame, Christophe Boëte

## Abstract

We report the complete CDS of five viruses: an iflavirus, a protoambidensovirus and three microviruses),which were detected by viromics surveillance of mosquitoes (*Aedes albopictus* and *Culex quinquefasciatus*) from the Réunion Island. We detected the protoambidensovirus, which belongs to aclade previously reported only in *C. pipiens*, in *A. albopictus*.

## Announcement

Our knowledge of mosquito viruses remains limited, although they can modulate the transmission of arboviruses **(1)**. We processed 13 *Aedes albopictus* and 9 *Culex quinquefasciatus* larvae samples collected in La Réunion Island in 2019. Morphological characteristics were used for species identification, and larvae were pooled by location. Samples weighted between 0.5g and 1.49g, and stored at −80°C. Viromes were obtained as described in **(2)**. We followed manufacturers’ instructions and default parameters except where otherwise noted. Samples were homogenised by a bead-beater,filtered through a 0.45 µm filter and centrifuged at 140,000 g for 2.5 hours. The pellets were resuspended, digested by DNaseI and RNaseA incubation at 37°C for 1.5 h. RNA and DNA were extracted using a NucleoSpin kit. Reverse transcription was performed by SuperScript III, cDNAs were purified by aQIAquick PCR Purification Kit and dsDNA synthetised by Klenow DNA polymerase I. DNA was amplified by random PCR, and PCR products were purified using a NucleoSpin gel and PCR clean-up kit. Libraries were prepared using a NEBNext Ultra DNA PCR free with Ilumina adapter kit without fragmentation step, and sequenced on an Illumina HiSeq3000 in 2×150bp paired-end mode. Adaptors were removed and reads were filtered for quality (q30 quality and read length >45 nt) using cutadapt 2.19 **(3)**. 10,150,317 reads were assembled into contigs by MEGAHIT 1.2.9 **(4)**. Taxonomic assignment was achieved using DIAMOND 0.9.30 with against the NCBI nr protein database **(5)**. Genome coverage was assessed by mapping using Bowtie2 3.5.1 (end™to-end very-sensitive) **(6)**. Open Reading Frames (ORFs) were identified using ORF finder (length cutoff >400 nt) on Geneious prime 2024.0.5 **(7)**, and were annotated by blastn query-centered alignment against RefSeq viral database on October 2024. Genome circularisation was performed using <u>Simple-Circularise</u> (overlap >20nt).

We reconstructed the complete circular genomes of three bacteriophages species from the *Microviridae* family (*Gokushovirinae* subfamily) (<u>PQ041302</u>, <u>PQ041303</u> and <u>PQ041304</u>). They share 54.5% to 75.0% of amino acid identity in major capsid and replication-associated proteins with their closest relatives, which were isolated from water environments **(Table 1)**. There is currently no species demarcation criteria based on genome similarity defined by the ICTV for the *Microviridae* family **(8)**. Their high abundance in *Aedes* samples suggests than they may infect its microbiota, although their presence could also be due to environmental or diet contamination.

**Table 1.** Information on the five viruses reconstructed from mosquito viromic data.

| Virus | GenBank<br>accession number | Genome |  | Coverage |  |  | Putative proteins |  |  | Closest identified relatives |  |  |  |
| --- | --- | --- | --- | --- | --- | --- | --- | --- | --- | --- | --- | --- | --- |
|  |  | size (nt) | %GC | average | number of reads | sample | name | size (nt) | size (aa) | virus name | accession<br>number | pairwise<br>identity (aa) | source |
| Culex quinquefasciatus associated densovirus, isolate 2019_VP12-D87 | PQ041300 | 5 459 | 36,7 | 12 521 | 501 573 | Cqfts virome sample 3 (D87)<br>SRR29133503 | NS1 | 687 | 229 | Culex pipiens densovirus | FJ810126 | 94,7 | Culex pipiens |
|  |  |  |  |  |  |  | NS2 | 372 | 214 |  |  | 92,2 |  |
|  |  |  |  |  |  |  | NS3 | 642 | 124 |  |  | 42,4 |  |
|  |  |  |  |  |  |  | VP | 2 208 | 736 |  |  | 90,5 |  |
| Culex quinquefasciatus associated iflavirus, isolate 2019_VP12-D85 | PQ041301 | 10 074 | 36,9 | 2 777 | 194 020 | Cqfts virome sample 1 (D85)<br>SRR29133486 | polyprotein | 9 231 | 3 077 | XiangYun picorna-like virus 4 | OL700176 | 98,5 | Culex theileri |
| Aedes albopictus associated microvirus 1, isolate 2019_VP12-D83 | PQ041302 | 4 365 | 43,9 | 243 | 8 198 | Aedes virome sample 11 (D83)<br>SRR29133492 | major capsid protein | 1 650 | 550 | Microvirus sp. isolate BS1_474 | MT309944 | 75,0 | wastewater |
|  |  |  |  |  |  |  | DNA pilot protein | 678 | 226 | Microviridae sp. isolate CD-SXD-WGA-1-k141_518849 | OR998997 | 43,3 | bacteria |
|  |  |  |  |  |  |  | internal scaffolding protein | 429 | 143 | Microvirus sp. isolate BS1_462 | MT309947 | 54,2 | wastewater |
|  |  |  |  |  |  |  | replication initiator protein | 1 023 | 341 | Arizlama microvirus | MW697615 | 54,9 | fresh water |
| Aedes albopictus associated microvirus 2, isolate 2019_VP12-D75 | PQ041303 | 4 387 | 46,3 | 62 | 2 200 | Aedes virome sample 3 (D75)<br>SRR29133501 | major capsid protein | 1 635 | 545 | Microviridae sp. isolate SD_MC_72 | MH572427 | 67,3 | Ciona robusta |
|  |  |  |  |  |  |  | DNA pilot protein | 801 | 267 | Microvirus sp. isolate LSGA_102419 | OR349779 | 23,4 | mollusc |
|  |  |  |  |  |  |  | internal scaffolding protein | 474 | 158 | Microviridae sp. isolate ctda862 | MH616902 | 65,8 | fish |
|  |  |  |  |  |  |  | replication initiator protein | 747 | 249 | Microvirus sp. isolate BS1_438 | MT309954 | 74,7 | wastewater |
| Aedes albopictus associated microvirus 3, isolate 2019_VP12-D82 | PQ041304 | 4 813 | 50,5 | 197 | 6 618 | Aedes virome sample 10 (D82)<br>SRR29133485 | major capsid protein | 1 653 | 551 | Microviridae sp. isolate ctcc708 | MH617066 | 73,4 | fish |
|  |  |  |  |  |  |  | DNA pilot protein | 795 | 265 | Microviridae sp. isolate ctji855 | MH617641 | 37,6 | fish |
|  |  |  |  |  |  |  | internal scaffolding protein | 444 | 148 | Microviridae sp. isolate ctcc708 | MH617066 | 61,9 | fish |
|  |  |  |  |  |  |  | replication initiator protein | 783 | 261 | Microvirus sp. isolate 1712115_622 | MT310309 | 68,6 | wastewater |

We report a complete CDS of a lineage of *XiangYun picorna-like virus 4* species (*Iflaviridae, Iflavirus*) (**Table 1**, <u>PQ041301</u>). It shares a whole polyprotein pairwise identity of 98.5% with a lineage discovered in *Culex theileri* from Yunnan, China (<u>OL700176</u>) **(9)**. This lineage clustered in a monophyletic clade containing only viral taxa isolated from dipteran species (mosquitoes and true flies), indicating that it is likely specific of dipterans (10.6084/m9.figshare.28565888). Further work is needed to determine the prevalence of XiangYun picorna-like virus 4 in natural populations of mosquitoes and its impact on mosquito fitness.

We report the complete CDS of a distant lineage of the mosquito infecting *Dipteran protoambidensovirus 1* species (*Parvoviridae, Densovirinae, Protoambidensovirus*, **Figure 1**, <u>PQ041300</u>).

**Figure 1.**
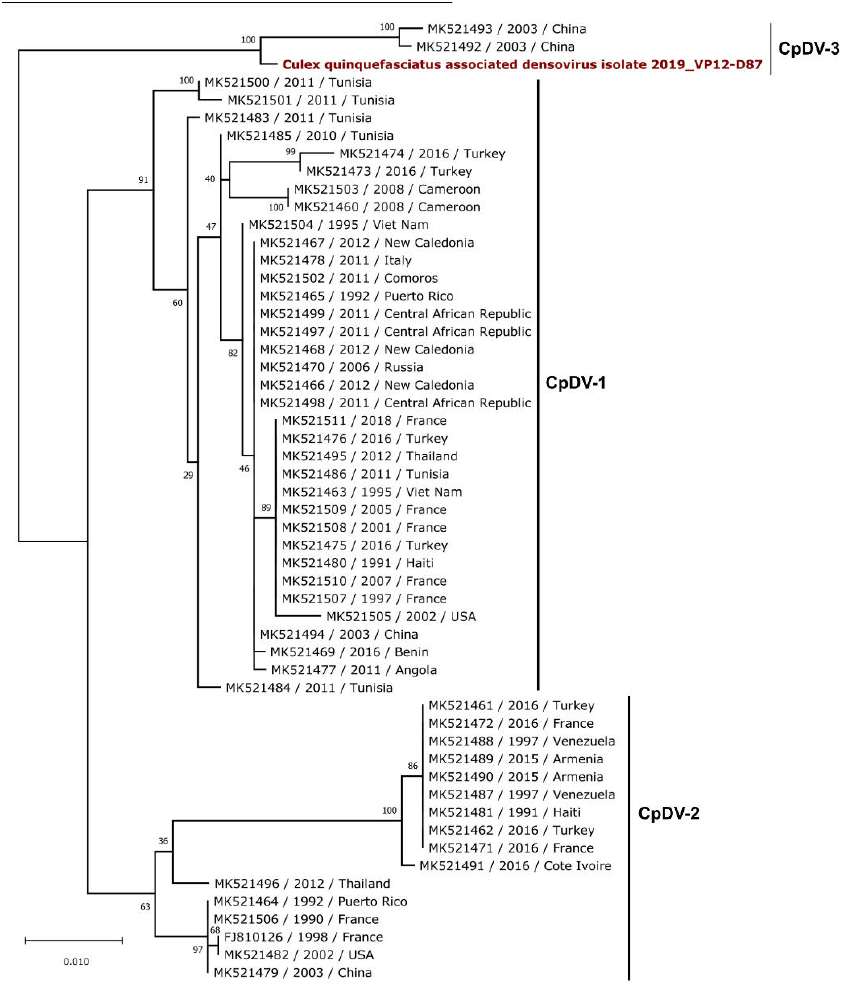
Maximum likelihood phylogenetic tree based on the NS1 nucleotide sequences of 52 Dipteran protoambidensovirus 1 (common virus name: Culex pipiens densovirus (CpDV)) sequences. The alignment of 892 nucleotides in length was produced by MAFFT v7.450 using the G-INS-i algorithm. The tree is mid-point rooted. Bootstrap values (100 replicates) are indicated at each node. Scale bar corresponds to nucleotide substitutions per site. The CpDV sequence obtained from our sample is coloured in red.

It shares 89.0% genome-wide nucleotide identity to its closest relative, Culex pipiens densovirus (<u>FJ810126.1</u>) **(Table 1) (10)**. This lineage belongs to the CpDV-3 clade which was previously only represented by sequences collected from Beijing (China) samples in 2003 **(11)**. Reported CpDV host range currently only includes *Culex pipiens* **(12,13)**, its broadening to *A. albopictus* warrants further confirmation.

## Data availability

The genomic sequences of the five full-length viral genomes or CDS have been deposited at GenBank under the accession numbers <u>PQ041300</u> to <u>PQ041304</u>. High-throughput sequencing reads were deposited in SRA under the accession no. <u>SRR29133481</u> to <u>SRR29133504</u> under <u>PRJNA1114772</u>BioProject.

## Acknowledgments

We thank Yann Gomard, Cyrille Lebon and Sarah Hafsah for their help in collecting samples in the field.

This work was funded by the French ANSES PNR EST programme (project 2018/1/183, “DENSOTOOL”, 2019–2023).

